# STING deficiency extends median lifespan and mitigates age-related liver pathology in mice

**DOI:** 10.64898/2026.10.16.759663

**Authors:** John C. Martinez, Sei Joong Kim, Joseph Cutting, John M. Sedivy, Andrei Seluanov, Vera Gorbunova

## Abstract

The cGAS-STING pathway is a major driver of chronic inflammation. While acute STING inhibition can reduce neuroinflammation and tissue damage, its long-term impact on longevity remains unknown. Here, we demonstrate that genetic deletion of STING extends median lifespan in mice. This survival benefit is accompanied by a marked resistance to age-related hepatic decline. Our findings demonstrate that targeting STING is a promising therapeutic target to extend healthy lifespan.

---

A crucial component of innate immunity in cells is the ability to respond to foreign genetic material invading the cytosol to avoid propagated infection. Double-stranded nucleic acids in the cytosol are recognized by cGAS, which then generates cGAMP and binds to STING to initiate interferon signaling in response^1,2,3^. The nature of cGAS-STING signaling relies solely on the presence of DNA in the cytoplasm; the inflammatory response is not biased towards viral DNA versus self-DNA. During cellular aging, DNA from mitochondria, LINE1s, and cytosolic chromatin fragments (CCFs) have been shown to active cGAS-STING signaling and drive inflammaging^4,5^. Targeting of cGAS-STING activity has shown promise in alleviating neurodegeneration by reducing inflammation^6^. Unexpectedly, we showed previously that cGAS deficient mice exhibit shortened lifespan, increased inflammation, and accelerates tissue damage and fibrosis^7^. However, this is likely due to an important, non-canonical function of cGAS in maintaining chromatin landscape in the nucleus as opposed to its role in type-I interferon production. Thus, alternative targeting of the pathway through cGAMP production or STING may have the potential to extended lifespan, healthspan, or both without undesirable dysregulation of nuclear chromatin. Here, we show that STING deficient mice display an extended median lifespan associated with significant reduction of age-related liver pathology and inflammation.

Unlike cGAS KO mice, STING KO mice display a nearly 10% extension in median lifespan relative to wild-type animals (**Fig. 1a**). At 25 months, STING KO mice showed a trend towards reduced frailty, scored using by Hewlett frailty index^8^ (**Fig. 1b**). Intriguingly, like cGAS KO, both sexes of STING KO mice maintain increased weight across life relative to WT with females specifically exhibiting increased body mass late in life (**Ex. Data 1a, b**). However, STING KO mice exhibit reduced hair greying with age. Hair greying has been linked to increased immune signaling within hair follicles and surrounding cells (**Fig. 1c**)^9,10^.

**Figure 1:**
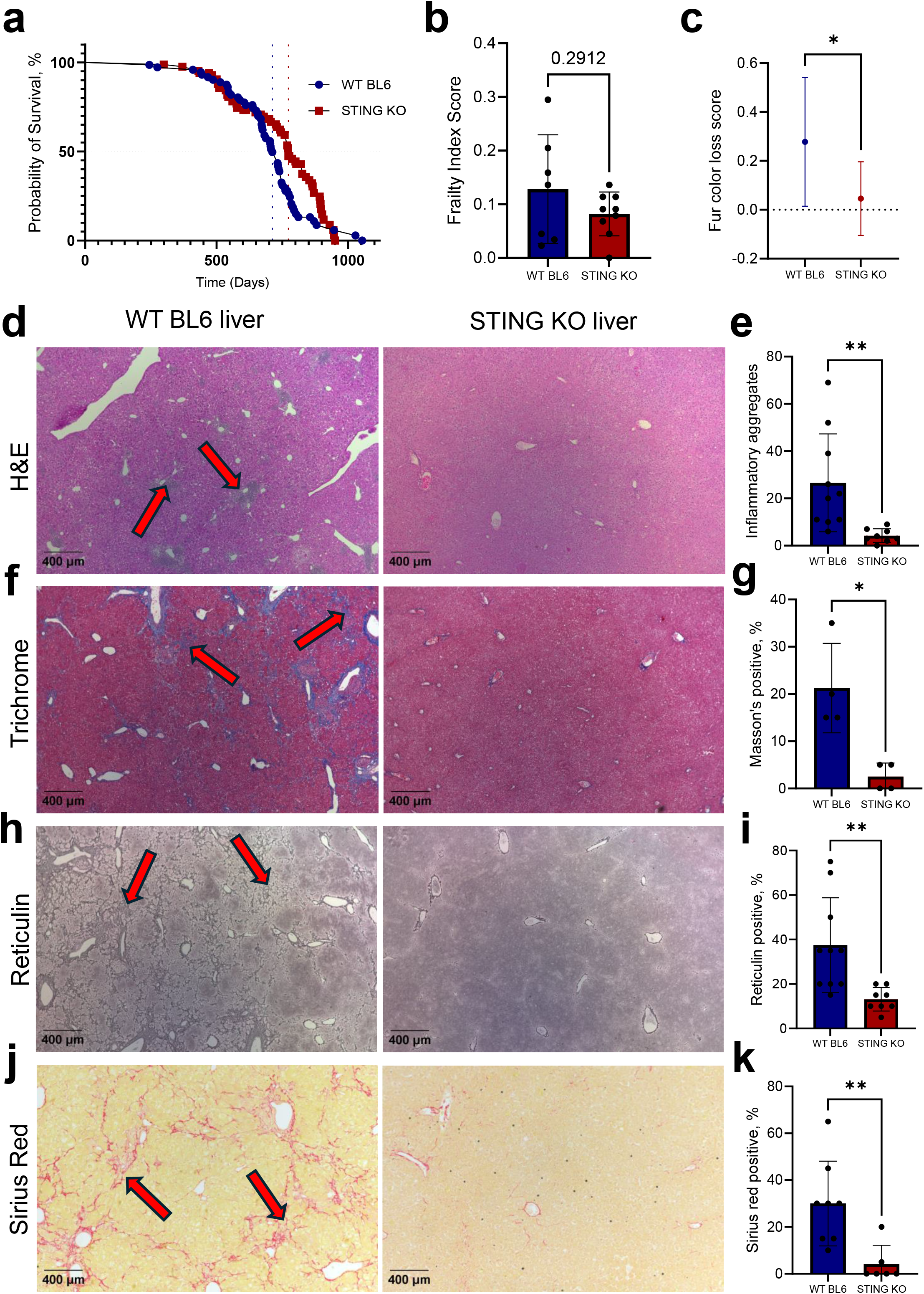
STING KO mice display extended median lifespan and resistance to age-associated liver damage. Survival of WT and STING KO mice. STING KO mice display a significantly extended median lifespan relative to WT mice (WT, n = 61, STING KO, n = 62; WT median survival = 712 days, STING KO median survival = 773 days). The Gehan-Breslow-Wilcoxon test was used (p = 0.0349). Blue and red dashed lines indicate median survival timepoint for WT BL6 and STING KO cohorts, respectively. Frailty index scoring of WT and STING KO male mice at 25-months-old (WT, n = 7, STING KO, n = 9; p = 0.291). Two-sided Welch’s t-test. Median fur color loss score for WT and STING KO male mice at 25-months-old. Mice are scored from 0 to 1 depending on severity of fur color loss (WT, n = 9, STING KO, n = 11; p = 0.0365). Two-sided Welch’s t-test. H&E staining of WT and STING KO male livers at 25-months-old. Relative to WT, STING KO livers exhibit significantly reduced inflammatory aggregate accumulation. Arrows indicate inflammatory aggregates. Quantification of inflammatory aggregate accumulation in livers (WT, n = 10, STING KO, n = 8; p = 0.0076). Two-sided Welch’s t-test. Masson’s-Trichrome staining of WT and STING KO male livers. STING KO livers exhibit reduced fibrosis with age. Arrows indicate Masson’s positive regions. Quantification of Masson’s-Trichrome-positive area of the liver from WT and STING KO mice (WT, n = 4; STING KO, n = 4; p = 0.0239). Two-sided Welch’s t-test. Reticulin staining of WT and STING KO male livers. STING KO mice exhibit reduced accumulation of early fibrotic fibers with age. Arrows indicate reticulin positive regions. Quantification of Reticulin-positive area of the liver from WT and STING KO mice (WT, n = 4, STING KO, n = 4; p = 0.0797). Two-sided Welch’s t-test. Sirius red staining of WT and STING KO male livers. STING KO mice exhibit reduced fibrosis with age. Arrows indicate Sirius red positive regions. Quantification of Sirius red-positive area of the liver from WT and STING KO mice (WT, n = 4, STING KO, n = 4; p = 0.049). Two-sided Welch’s t-test.

Histological examination by H&E staining of heart, lung, liver, kidney, and pancreas revealed negligible differences across all tissues except liver, which displayed a strong reduction in inflammatory aggregates in the STING KO (**Fig. 1d, e**). Masson’s trichrome, reticulin, and Sirius red staining showed a striking reduction in fibrosis in STING KO livers in both sexes (**Fig. 1f-k; Ex. Data 1c**). cGAS-STING signaling has been proposed as a potential therapeutic target for attenuating liver damage and has been shown to be active and upregulated during liver fibrosis^11,12,13,14,15^. STING specifically has been shown to be activated during both alcoholic and non-alcoholic fatty liver disease^16,17,18,19^. Comparatively, we and others have demonstrated that cGAS KO accelerates liver damage rather than alleviate it^20^. Crucially, it has also been demonstrated that STING KO mice are more resistant to liver damage induced by acetaminophen overdose comparted to WT animals^21^. Intriguingly, STING mediated liver damage appears to be dependent on activation of the NLRP3 inflammasome complex formation rather than canonical cGAS-STING signaling^14,22^. Our study demonstrates that STING KO is also able to resist general age-associated liver damage.

We next analyzed the transcriptomes of STING KO and WT livers. We performed RNA-seq on the livers of young (12-months) and old (25-months) WT and STING KO male mice. We observed a large number of differentially expressed genes in the aged mice, with the KO exhibiting 907 significantly down regulated genes versus 240 upregulated genes (**Fig. 2a**). Comparing mRNA levels between young and old WT livers revealed STING as significantly elevated in aged livers **(Ex. Data 2)**. Analysis of specific genes from the RNAseq confirmed significant reduction in expression of fibrotic and inflammatory genes (**Fig. 2b**). GO term analysis further demonstrated that some of the most significantly downregulated pathways in aged STING KO liver are related to inflammation (AIM2 inflammasome assembly, leukocyte migration, cytokine production, interferon-mediated signaling) and fibrosis related (extracellular matrix organization, collagen metabolic process, tissue remodeling) (**Fig. 2c**). Employing RT-qPCR on mouse livers confirmed that aged STING KO mice are protected from age-associated activation of fibrosis genes including *Col1a1, Col3a1, TMP-1*, and *αSMA* (**Fig. 2d; Ex. Data 3a**). Probing for inflammation, we observed that STING KO livers also resist activation of *IL-6, CD68*, and *IL-1β* (**Ex. Data 3a**). Finally, we performed immunohistochemistry (IHC) on livers from middle-aged (18-months) and aged (25-months) WT livers for activated, phosphorylated-STING **(Fig. 2e; Ex. Data 3b)**. While mice at 18-months retain low levels of phosphorylated-STING, 25-month-old livers displayed strong, systematic increase in phosphorylated-STING. These results show that activated phosphorylated-STING in the liver increases significantly with age. Despite cGAS and STING working together in the cytosolic DNA sensing pathway, their knockouts have dramatically different effects on aging. cGAS knockout shortens lifespan and increases inflammation due to loss of the non-canonical function of cGAS in maintaining chromatin landscape, while STING knockout extends lifespan and attenuates age-related inflammation^7^. Taken together, our results indicate that targeting STING, rather than cGAS, has the potential to attenuate age-associated liver damage, inflammation, and extend lifespan.

**Figure 2:**
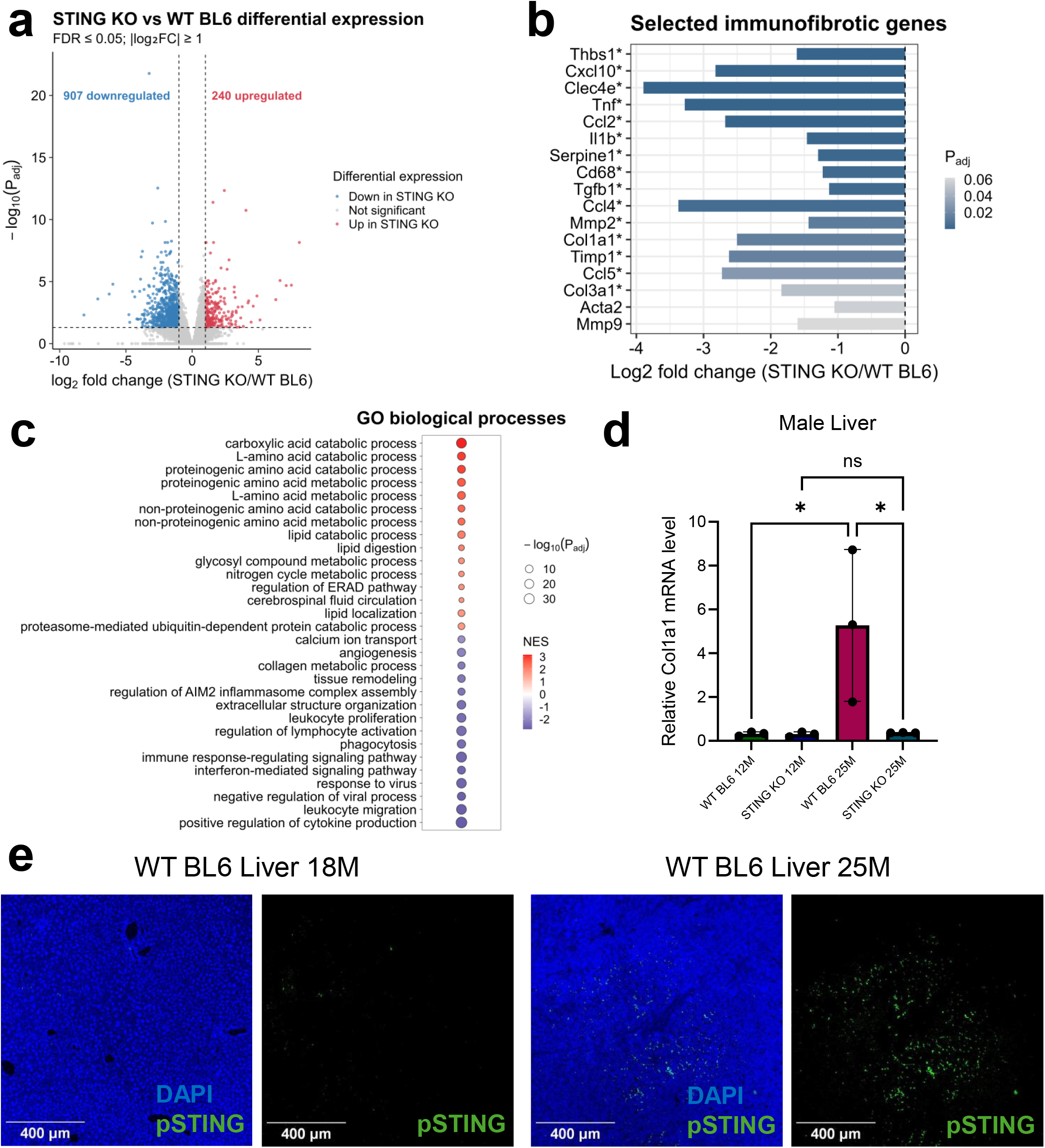
Livers from STING KO mice show reduced pro-inflammatory and fibrotic expression signatures. Volcano plot displaying differentially expressed genes from WT and STING KO male liver tissue at 25-months-old. Relative to WT, STING KO livers display 240 significantly upregulated genes and 907 downregulated genes. Differential expression was assessed using the DESeq2 Wald test with Benjamini-Hochberg adjustment, with significance defined as adjusted P ≤ 0.05 and |log2 fold change| ≥ 1 (WT, n = 3; STING KO, n = 3). Expression of selected inflammatory and fibrotic genes in 25-month-old STING KO livers relative to WT. Asterisks indicate adjusted P < 0.05. Top biological processes gene ontology (GO) terms for aged STING KO versus WT liver tissue showing repression of pro-inflammatory and fibrotic pathways in STING KO. RT-qPCR confirming increased Col1a1 mRNA levels in aged WT, but not aged STING KO, livers (n = 3 for each group; p = 0.0334 between 12-month and 25-month WT; p = 0.0349 between 25-month WT and STING KO). One-way ANOVA test. Representative immunohistochemistry (IHC) staining for pSTING in 18- and 25-month-old WT livers. pSTING accumulates in WT mouse livers with age.

## Materials and Methods

### Animal Care and Pathogen Testing

STING KO and wild-type mice were obtained from Jackson Laboratories (strain B6-Sting^tm1.2Camb^/J, #025805) and cared for in accordance with the regulations developed and approved by the University of Rochester Committee on Animal Resources (UCAR).^23^ UCAR is compliant with FDA and NIH (National Institutes of Health) animal care guidelines and reviews all protocols before approval and implementation (#2017-027). Animals were housed in a one-way facility utilizing microisolator technology. Frailty scoring conducted according to established protocol^8^.

Triennial health surveillance is performed using PCR testing of exhaust air from the ventilated racks. Screening if performed for Mouse Rotavirus (EDIM), Theiler’s Murine Encephalomyelitis Virus (TMEV), Mouse Hepatitis Virus (MHV), Mycoplasma pulmonis, Mouse Parvovirus (MPV), Minute Virus of Mice (MMV), Pneumonia Virus of Mice (PVM), Sendai Virus, Helicobacter, pinworms and fur mites during two of these testing periods. In the third period, testing is expanded to also include CAR bacillus, Lymphocytic Choriomeningitis Virus (LCMV), Ectromelia Virus, Mouse Adenovirus (MAV), Mouse Cytomegalovirus (MCMV) and Reovirus.

### Tissue histology, hematoxylin & eosin (H&E), Masson’s trichrome, reticulin, and Sirius red staining, and imaging

Mice were sacrificed via isoflurane exposure and perfused with PBS. Tissues were then isolated from mice, placed in a histology cassette, and fixed in 4% paraformaldehyde (PFA) for 48 hours. Tissues were subsequently submitted to the University of Rochester CMSR Histology Core (NIH P30 AR069655) for paraffin embedment, sectioning, and staining (H&E, Masson’s, and reticulin). Sirius red staining was performed using kit from ABCAM (ab150681) following manufacturer’s protocol. Slides were imaged using brightfield microscopy.

### Immunohistochemistry (IHC) for STING and pSTING

Livers were isolated from WT and STING KO mice at 18 and 27 months old and processed by the University of Rochester Histology Core as described above. Tissue sections were then rehydrated with two xylene washes at 10-minutes each followed by 2-minute washes in 100% EtOH (2X), 90% EtOH, 75% EtOH, and 50% EtOH. Slides were briefly washed in PBS then rinsed in running water for 3-minutes. Next, tissues were blocked in PBS with 4% BSA, 2% donkey serum, and 0.1% Triton-X-100 for 1-hour. Sections were stained with pSTING (Cell Signaling #72971) at a 1:200 dilution in a humidity chamber overnight at 4-degrees Celsius. The next day, slides were washed 3 times with 0.1% Triton-X-100 in PBS before being stained with Alexa-488 (Invitrogen #A32731) at a 1:1000 dilution for 2-hours at room temperature. Slides were then washed an additional 3 times with 0.1% Triton-X-100 before being stained with 2µg/mL DAPI (Thermo-Fisher #62248) in 0.1% Triton-X-100 for 15-minutes. Finally, slides were mounted in Permount medium (Fisher Chemical #SP15-100) and stored at 4-degrees Celsius until imaged.

### Tissue processing, RNA extraction, and RNAseq library preparation

Tissues were isolated from mice, flash frozen in liquid nitrogen, then pulverized using piton/cell crusher chilled using liquid nitrogen to a fine powder. RNA was isolated from pulverized tissue using Trizol. Insoluble material was removed by spinning tubes for 1 minute at 12,000 g/4-degrees and transferring supernatant to new tube. 100 µl chloroform was added to this solution, shaken vigorously for 15 seconds, incubated at room temperature for 5 minutes, then centrifuged for 15 minutes at 12,000 g. Aqueous phase containing RNA was then extracted to a new tube, mixed with 250 µl isopropanol, then incubated at room temperature for 10 minutes. The samples were then centrifuged for 10 minutes at 12,000 g. Supernatant was then removed, leaving behind RNA pellet. The pellet was then washed in 500 µl 75% ethanol in DPEC ddH2O and centrifuged at 8,000 g for 5 minutes. EtOH was then removed, and the pellet allowed to air dry for 10 minutes. RNA was further purified utilizing RNA PureLink Mini Kit (Thermo). Libraries for RNAseq were generated using NEBNext^®^ Ultra^™^ II Directional RNA Library Prep Kit for Illumina^®^ from New England Biolabs following manufacturer’s protocol (E7760).

### Quantitative RT-PCR

RNA for RT-qPCR was isolated and purified using method above. cDNA was then generated using iScript cDNA synthesis kit from BioRad (1708890) using random hexamer primers. qRT-PCR was performed using the Bio-Rad CFX Connect Real Time machine and SYBR Green Master Mix (Bio-Rad) using 30 ng/cDNA per reaction. RNA expression was normalized to EEF2. Primers utilized: EEF2 FWD: GATGATCACCATCCACTTACC; EEF2 REV: GGGTCGCAGCTCTTAATAC; Col1a1 FWD: TGACTGGAAGAGCGGAGAGATCT; Col1a1 REV: TTCGGGCTGATGTACCAGTTC; Col3a1 FWD: CCTTCTACACCTGCTCCT; Col3a1 REV: CTTCCTGACTCTCCATCCT; αSMA FWD: TGCCGAGCGTGAGATTGTC; αSMA REV: CGTTCGTTTCCAATGGTGATC; TMP-1 FWD: CCAGAACCGCAGTGAAGAGT; TMP-1 REV: TCTGGTAGTCCTCAGAGCCC; CD68 FWD: GGGGCTCTTGGGAACTACAC; CD68 REV: GTACCGTCACAACCTCCCTG; IL-6 FWD: TACCACTTCACAAGTCGGAGGC; IL-6 REV: CTGCAAGTGCATCATCGTTGTTC; IL-1β FWD: CCGTGGACCTTCCAGGATGA; IL-1β REV: GGGAACGTCACACACCAGCA.

## Quantification and Statistical Analysis

Unless specified otherwise, the student’s t-test was utilized for comparison between groups. All tests were two-tailed, and p-values were considered significant below the 0.05 threshold.

### RNAseq analysis

Paired-end RNA-seq reads were adapter- and quality-trimmed using Trim Galore v0.6.2 and aligned to the GRCm39 mouse genome with HISAT2 v2.1.0. The resulting alignments were converted to sorted and indexed BAM files using SAMtools 1.21. Gene-level counts were generated with featureCounts v2.0.3 using the GRCm39 gene annotation in unstranded, paired-end mode. Count matrices were analyzed in R with DESeq2 v1.48.1. Genes with counts ≥10 in at least three samples were retained, and differential expression was analyzed using a design formula of ∼ age * genotype, with the KO versus WT effect at 25 months extracted as a model contrast. Differentially expressed genes were defined as those with Benjamini-Hochberg-adjusted P ≤ 0.05 and |log2 fold change| ≥ 1. Gene Ontology biological process enrichment analysis was performed using clusterProfiler v4.16.0 with org.Mm.eg.db v3.21.0 annotations, using genes ranked by the DESeq2 Wald statistic. Redundant GO terms were reduced using simplify() from clusterProfiler with a similarity cutoff of 0.5, and the top 20 terms were displayed^24,25^.

## Code Availability

Code used for data processing, statistical analysis, and figure generation will be made publicly available upon publication.

## Acknowledgements

We would like to thank all members of V.G.’s and A.S.’s laboratories for their helpful discussions.

## Author Contributions

J.M., A.S., and V.G. conceived the experiments. J.M., S.J.K, and J.C. performed the experiments. J.M., A.S., and V.G. analyzed the data. S.J.K. conducted bioinformatic analysis. J.S., A.S., and V.G. secured funding for various aspects of the project. J.M., A.S., and V.G. wrote the manuscript with input from all authors.

## Figure Legends

**Extended Data Fig. 1:**
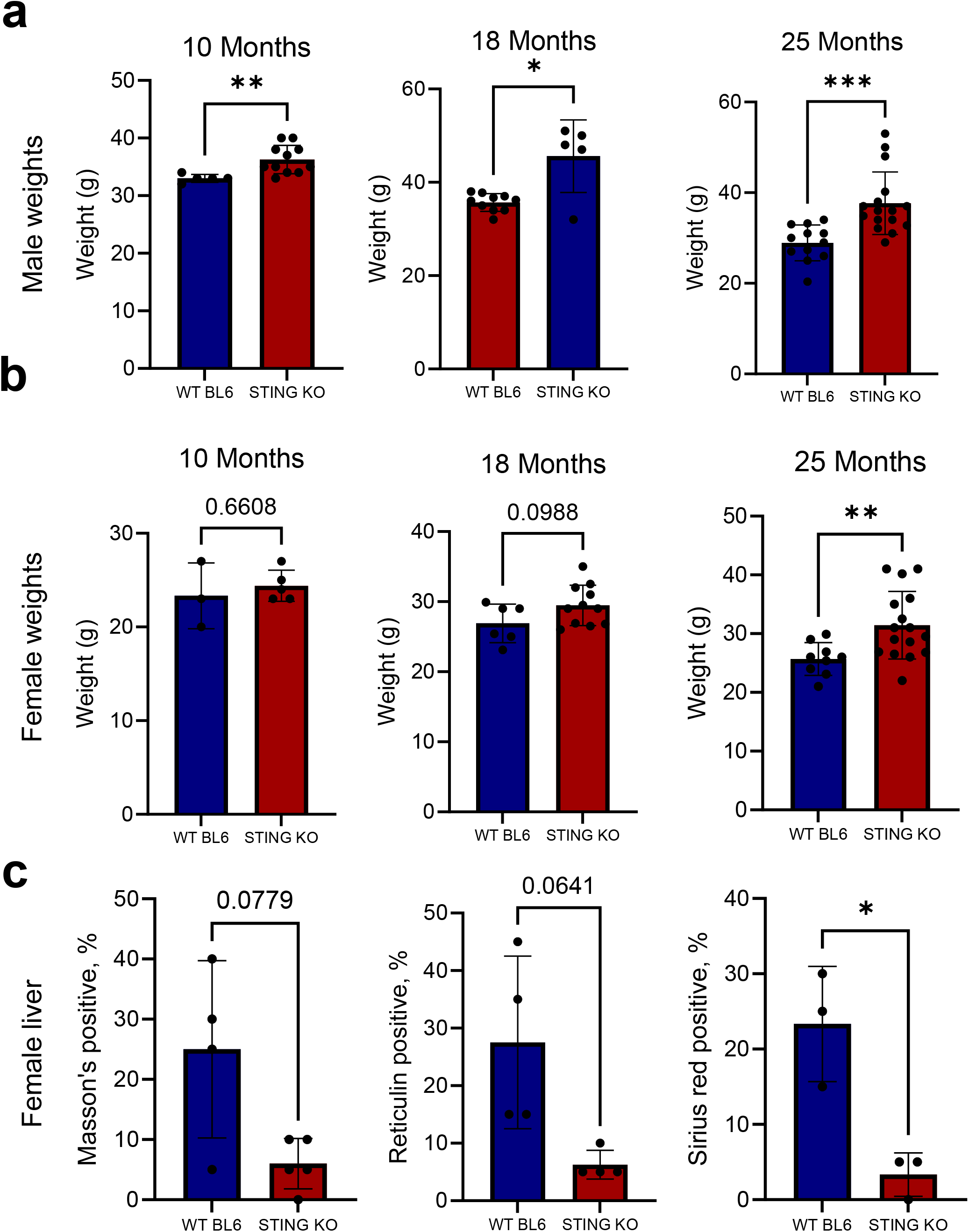
STING KO mice exhibit increased body weight but attenuated liver damage. Aged STING KO male mice exhibit increased body mass (10-month: WT, n = 5, STING KO, n = 11; p = 0.0013; 18-month: WT, n = 10, STING KO, n = 5; p = 0.0448; 25-month: WT, n = 12, STING KO, n = 16; p = 0.0003). Two-sided Welch’s t-test. STING KO female mice exhibit increased body mass late in life (10-month: WT, n = 3, STING KO, n = 5; p = 0.6608; 18-month: WT, n = 6, STING KO, n = 11; p = 0.0988; 25-month: WT, n = 9, STING KO, n = 16; p = 0.0028). Two-sided Welch’s t-test. Quantification of Masson’s-Trichrome-positive area of the liver from WT and STING KO female mice (WT, n = 4; STING KO, n = 5; p = 0.0779). Quantification of Reticulin-positive area of the liver from WT and STING KO female mice (WT, n = 4, STING KO, n = 4; p = 0.0641). Quantification of Sirius red-positive area of the liver from WT and STING KO female mice (WT, n = 3, STING KO, n = 3; p = 0.0327). All two-sided Welch’s t-test.

**Extended Data Fig. 2:**
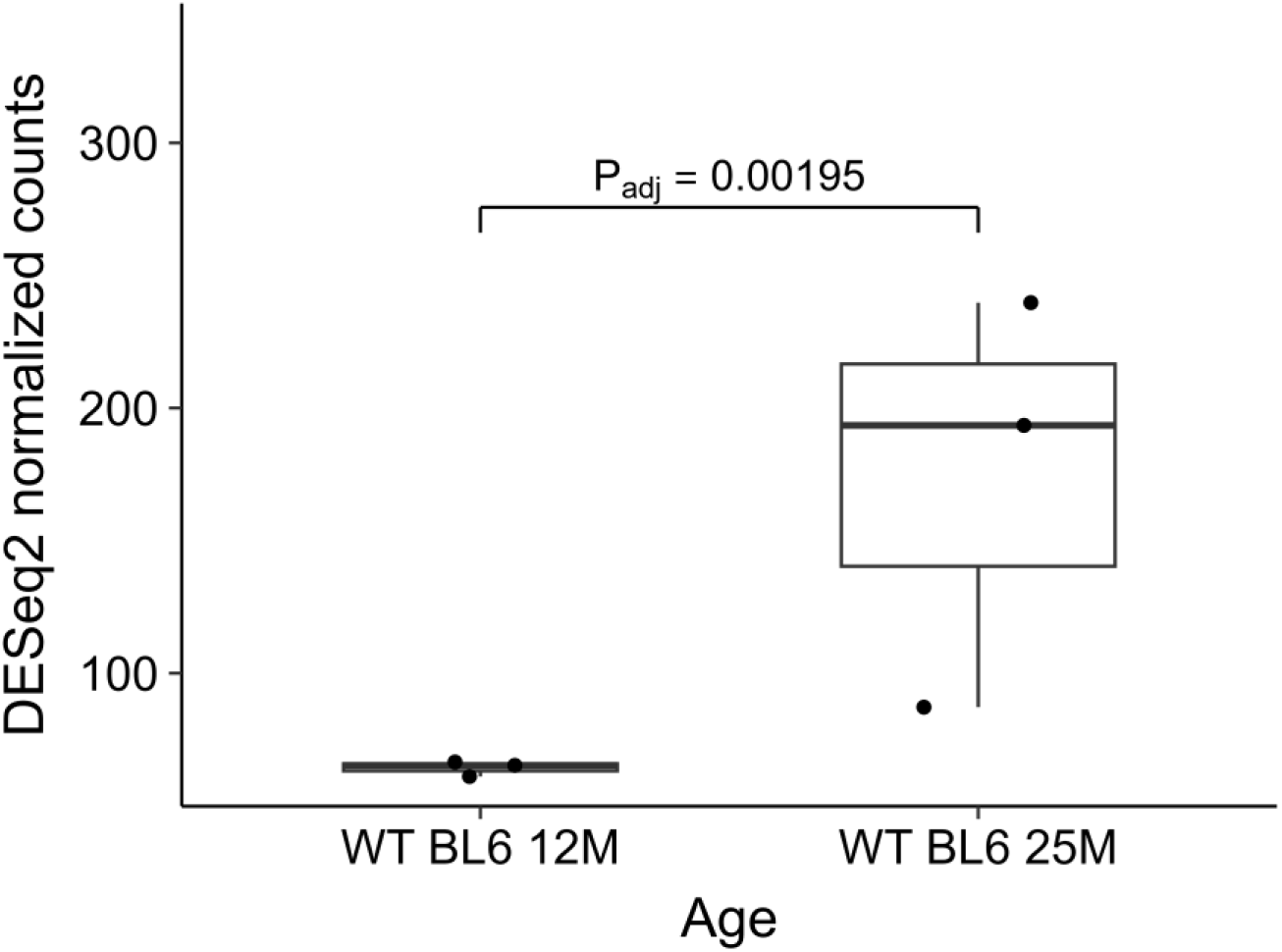
STING mRNA levels increase during aging. STING mRNA levels, measured by RNAseq, increase in aged WT mice (n = 3 per age group; adjusted P = 0.00195). DESeq2 normalized counts. Statistical significance was determined using the DESeq2 Wald test with Benjamini-Hochberg correction.

**Extended Data Fig. 3:**
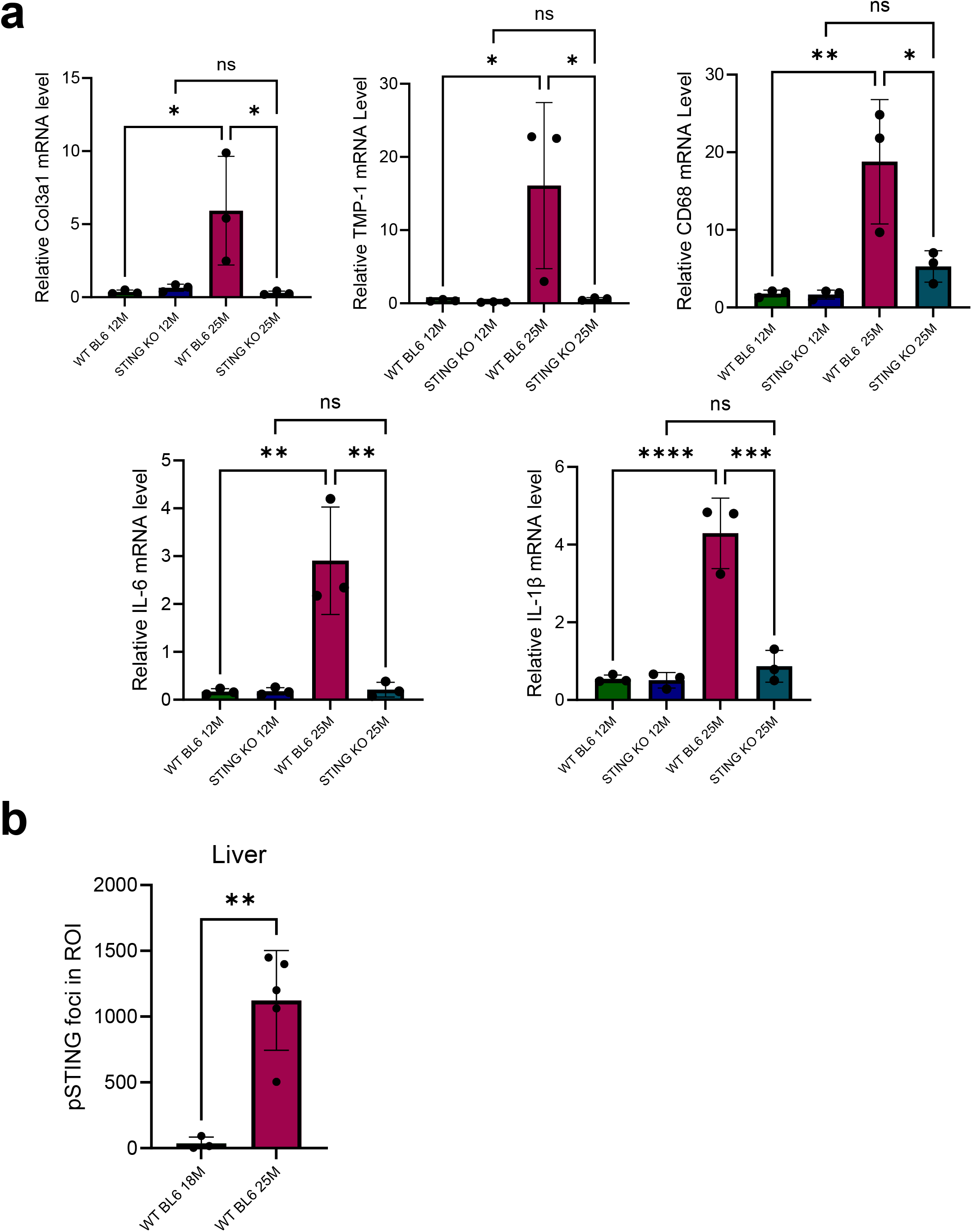
Aged STING KO livers are resistant to age-associated fibrosis. RT-qPCR confirming increased Col3a1, TMP-1, CD68, IL-6, and IL-1β mRNA levels in aged WT, but not aged STING KO, livers (n = 3 for each group; Col3a1: p = 0.0267 between 12-month and 25-month WT; p = 0.0252 between 25-month WT and STING KO; TMP-1: p = 0.0387 between 12-month and 25-month WT; p = 0.0412 between 25-month WT and STING KO; CD68: p = 0.0045 between 12-month and 25-month WT; p = 0.0170 between 25-month WT and STING KO; IL-6: p = 0.0016 between 12-month and 25-month WT; p = 0.0018 between 25-month WT and STING KO; IL-1β: p < 0.0001 between 12-month and 25-month WT; p = 0.0002 between 25-month WT and STING KO). All one-way ANOVA test. Quantification of pSTING staining in young and old WT livers. ImageJ was used to count pSTING+ cells in sample images (WT, n = 3, STING KO, n = 5; p = 0.0027). Two-sided Welch’s t-test.

